# Mechanism of action of rigosertib does not involve tubulin binding

**DOI:** 10.1101/2019.12.12.874719

**Authors:** Stacey J. Baker, Stephen C. Cosenza, Saikrishna Athuluri-Divakar, M.V. Ramana Reddy, Rodrigo Vasquez-Del Carpio, Rinku Jain, Aneel K Aggarwal, E. Premkumar Reddy

## Abstract

Rigosertib is a novel benzyl styryl sulfone that inhibits the growth of a wide variety of human tumor cells *in vitro* and *in vivo* and is currently in Phase III clinical trials. We recently provided structural and biochemical evidence to show that rigosertib acts as a RAS-mimetic by binding to Ras Binding Domains (RBDs) of the RAF and PI3K family proteins and disrupts their binding to RAS. In a recent study, Jost et al (2017) attributed the mechanism of action of rigosertib to microtubule-binding. In these studies, rigosertib was obtained from a commercial vendor. We have been unable to replicate the reported results with clinical grade rigosertib, and hence compared the purity of clinical grade and commercially sourced rigosertib. We find that the commercially sourced rigosertib contains approximately 5% ON01500, a potent inhibitor of tubulin polymerization. Clinical grade rigosertib, which is free of this impurity, does not exhibit tubulin binding activity. *In vivo*, cell lines that express mutant β-tubulin (TUBBL240F) were also reported to be resistant to the effects of rigosertib. However, our studies showed that both wild-type and TUBBL240F-expressing cells failed to proliferate in the presence of rigosertib at concentrations that are lethal to wild-type cells. Morphologically, we find that rigosertib, at lethal concentrations, induced a senescence-like phenotype in the small percentage of both wild-type and TUBBL240F-expressing cells that survive in the presence of rigosertib. Our results suggest that TUBBL240F expressing cells are more prone to undergo senescence in the presence of rigosertib as well as BI2536, an unrelated ATP-competitive pan-PLK inhibitor. The appearance of these senescent cells could be incorrectly scored as resistant cells in flow cytometric assays using short term cultures.

## IINTRODUCTION

Rigosertib is a novel benzyl styryl sulfone (figure 1A) that inhibits the growth of a wide variety of human tumor cells *in vitro* and impairs tumor growth *in vivo* with little toxicity (Jimeno et al., 2009; Reddy et al., 2011; Agoni et al., 2014; Silverman et al., 2014). We recently described the mechanism of action of rigosertib (Athuluri-Divakar et al, 2016) where we provided structural and biochemical evidence to show that rigosertib acts as a RAS-mimetic and binds to the RBDs of the RAF and PI3K family proteins and disrupts their ability to bind to RAS. This conclusion was based on a number of biochemical assays, which included chemical pulldown of target proteins by rigosertib bound to agarose beads, thermal shift assays, microscale thermophoresis (MST), as well and nuclear magnetic resonance (NMR) spectroscopy to derive the structure of the rigosertib-RAF-RBD complex. We also demonstrated that this compound inhibits RAS-mediated activation of MAPK and AKT pathways in *in vitro* as well as animal models of RAS-induced tumors.

**Figure 1.**
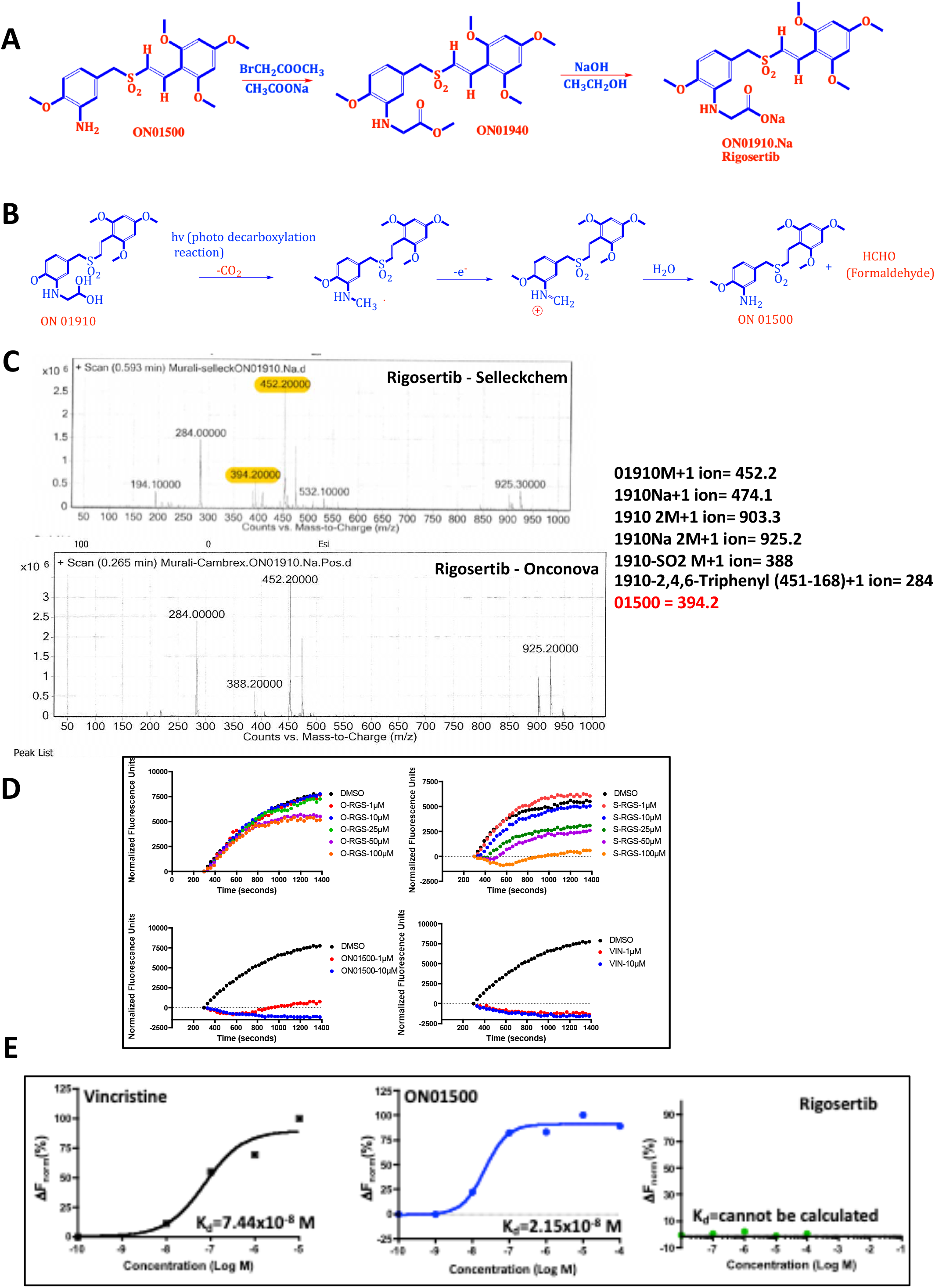
Purity and tubulin-binding activities of pharmaceutical grade and commercial grade rigosertib. **(A)** Synthetic scheme used for the preparation of rigosertib (Reddy et al, 2011). **(B)** Photo degradation of rigosertib to ON01500. **(C)** LC/MS/MS analysis of rigosertib obtained from Onconova Therapeutics Inc. and Selleckchem. The two compounds were analyzed on Agilent 6410 triple quadrupole mass spectrometer. Samples were dissolved in acetonitrile and eluted in 60% ammonium acetate and 40% acetonitrile on C18 column at a flow rate of 0.2 mL/min. The analytes were monitored by tandem mass spectrometry with electrospray positive ionization. **(D)** Polymerization of tubulin in the presence of rigosertib from Onconova Therapeutics (O-RGS), Selleckchem (S-RGS), ON01500 (Onconova Therapeutics) and vincristine (VIN). 25μg of MAP-rich tubulin in general tubulin buffer was mixed with 1mM GTP and fluorescence reporter in the presence of vehicle (DMSO) or increasing concentrations of the indicated compound. Tubulin polymerization as a function of fluorescence was recorded over the indicated time at 37°C. **(E)** Microscale thermophoretic analysis of ON01500 and vincristine with purified tubulin. Tubulin was labeled using the Monolith NT protein labeling kit RED-NHS according to the instructions of the manufacturer. Labeled protein was incubated with increasing concentrations of ON01500 or vincristine for 30 minutes and subjected to microscale thermophoresis.

In a recent study, Jost et al (2017) reported microtubule-binding activity of rigosertib, which was purchased from a commercial vendor. In the binding studies, Jost et al. reported that rigosertib, at high micromolar concentrations (>20μM), showed microtubule depolymerizing activity. Since we were unable to reproduce the results of Jost et al (2017), and impurities present in non-clinical grade preparations of rigosertib could have a significant effect on binding results, we examined the chemical and molecular basis for this discrepancy. We find that commercial grade preparations of rigosertib are often contaminated with synthetic intermediates and degradation products of rigosertib which contribute to the tubulin depolymerizing activity of these preparations. In addition, our results show that wild-type and TUBBL240F-expressing cells failed to proliferate in the presence of clinical grade rigosertib at concentrations that are lethal to wild-type cells. We also find that rigosertib, at lethal concentrations, induced a senescence-like phenotype in the small percentages of both wild-type and TUBBL240F-expressing cells and the appearance of these senescent cells could be incorrectly scored as resistant cells in flow cytometric assays using short term cultures.

## RESULTS

### Commercial Preparations of Rigosertib Are Often Contaminated With ON01500, an Intermediate With Potent Tubulin Depolymerizing Activity

A key late stage intermediate in the synthesis of rigosertib is ON01500 (figure 1A), which is an amino styryl-benzyl sulfone with potent tubulin-binding and depolymerization activity. Conversion of the amino group of ON01500 into a glycyl moiety results in the formation of ON01910 (rigosertib) (Reddy et al, 2011). Since the two compounds have distinct physiochemical properties (such as aqueous solubility), final purification and quality control measures are incorporated in the cGMP manufacturing of rigosertib, which is conducted under contract by the company (Onconova Therapeutics, Inc.) conducting clinical trials of this compound. A typical batch of clinical material is 99.9% pure and could include 0.1% of these impurities. Several storage conditions, including higher temperature, acidic pH and exposure to intense light can lead to the degradation of rigosertib into ON01500, as shown in figure 1B (Patel et al, 2017 and data not shown). We compared several batches of material obtained from the unlicensed vendor (Selleckchem) with materials obtained directly from Onconova Therapeutics.

To determine whether these preparations of rigosertib are contaminated with ON01500, we analyzed both preparations by NMR and mass spectrometry. The results of one such study are presented in figure 1C, and show that this particular batch of rigosertib purchased from Selleckchem contained approximately 5% of ON01500 as well as a few additional contaminants, while clinical grade rigosertib obtained from Onconova Therapeutics had undetectable amounts of these contaminants.

Next, we compared the tubulin depolymerizing activities of clinical grade rigosertib provided by Onconova Therapeutics (O-RGS), commercial grade rigosertib purchased from Selleckchem (S-RGS) and ON01500. Vincristine was used as a positive control. Tubulin polymerization assays were performed using MAP-rich tubulin and a fluorescence-based dye (Cytoskeleton Inc) that, when incorporated into polymerizing tubulin, results in fluorescence enhancement as polymerization occurs. The results of this study, presented in figure 1D, show that the highly purified preparation of rigosertib obtained from Onconova (O-RGS) shows little or no tubulin depolymerizing activity at doses up to 50μM. We observed a delay in tubulin polymerization at concentrations of 50μM and 100μM, which could be attributed to the contaminants which are less than 0.1%. These concentrations are ~500-1,000-fold higher than the GI50 value of rigosertib for any tumor cell line that is sensitive to the effects this compound. In contrast, tubulin was completely depolymerized in the presence of as little as 1μM ON01500 or vincristine. Rigosertib purchased from Selleckchem (S-RGS), on the other hand, exhibits depolymerizing activity when used at concentrations above 25μM, reaching complete depolymerization at a concentration of 100μM (Fig. 1D), suggestive of the effect of the impurity.

Next, we examined the binding affinity of pure ON01500, O-RGS and vincristine to purified tubulin preparations using microscale thermophoresis (MST) (Wienken et al., 2010). The results of this study, shown in figure 1E, demonstrate that both vincristine and ON01500 bind to tubulin with similar high affinities, which are reflected in their dissociation constants of 74 and 21nM, respectively. When we examined the binding of O-RGS using this technique, we were unable to detect binding even at a concentration of 50μM. While there was a hint of association at 100μM, the binding curve did not reach a plateau even at 1mM, suggesting that O-RGS does not have detectable tubulin-binding activity.

### Crystallography Studies Using Commercial Grade Rigosertib with ⇓–Tνβνλιν

Jost et al (2017) also report the crystal structure of tubulin in complex with commercially sourced rigosertib. Figure 2 shows the 2F_o_-F_c_ electron density for the modeled rigosertib (PDB Id: 5OV7) displayed at 0.5σ and 0.8σ, as well as at the more conventional 1σ cutoffs. The electron density for the carboxy group is relatively weak and the proposed hydrogen bonding interactions with αS178 (distances of 3.9 and 3.0Å for molecules B and D) and ⇓Lys352 (5.1 and 4.2 Å respectively) are less than optimal. In our opinion, a more likely interpretation of the weak electron density would be that a water molecule with the capacity to establish hydrogen bonding interactions with the side chain of βAsn349 is occupying this space. In light of our data showing the presence ON01500 in commercially sourced rigosertib, and the capacity of ON01500 to bind and depolymerize tubulins, the proposed structure is arguably more consistent with the binding of ON 01500 and a water molecule rather than rigosertib to tubulin.

**Figure 2.**
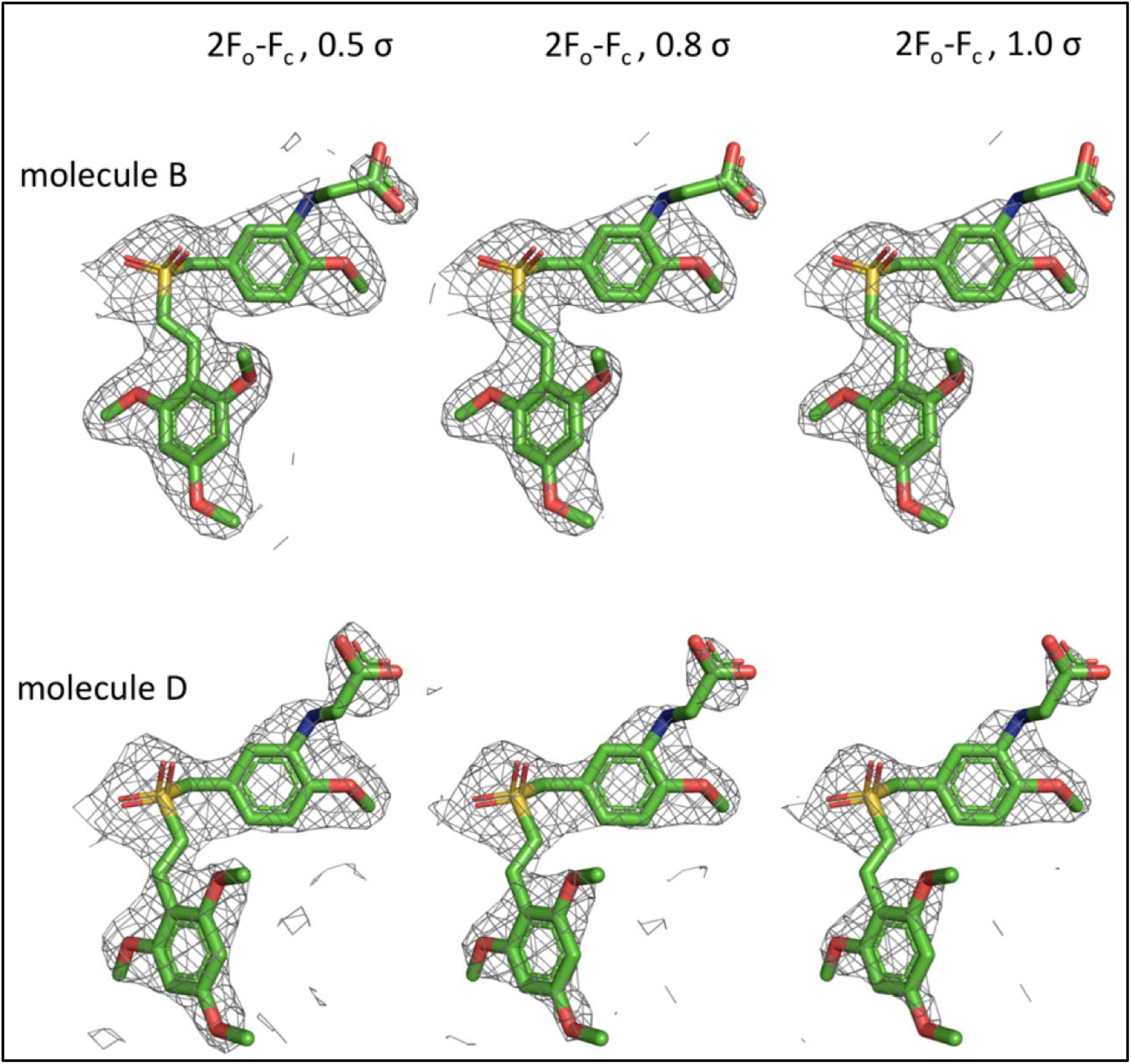
2F_o_-F_c_ electron density map (gray) for rigosertib modeled in the two αβ-tubulin heterodimers in the asymmetric unit of 5ov7. Density is contoured at 0.5σ (left), 0.8σ (middle) and at 1σ (right).

### L240F Tubulin Mutant Does Not Confer Resistance to Rigosertib

Jost et al. (2017) reported that expression of the L240F beta-tubulin mutant provides resistance to rigosertib, suggesting that tubulin binding is critical to its cytotoxic activity. For these experiments, Jost et al. (2017) prepared lentiviral constructs that encoded empty vector or wildtype (wt) *TUBB* or *TUBB* L240F, which were individually transduced into wt K562, HeLa, or H358 cells. These cell lines, which express the mCherry marker, were combined in a 1:1 ratio with their respective parental lines, treated with rigosertib or DMSO and the fraction of *TUBB*-expressing cells measured up to 7 days after treatment as mCherry-positive cells by flow cytometry. An elevated ratio of mCherry-expressing cells after rigosertib treatment compared to DMSO was interpreted to indicate that expression of the L240F mutant tubulin confers resistance. To examine this phenomenon in greater detail, we obtained lentiviral vectors that encode wt tubulin or the L240F beta-tubulin mutant from Dr. Weissman’s laboratory and repeated these studies using K562 cells that were transduced with empty vector or the TUBB L240F expression vector. Seventy-two hours post-infection, the cells were mixed 1:1 with uninfected cells and treated with DMSO, increasing concentrations of rigosertib or BI2536, a pan-PLK inhibitor as a control. Growth curves of the mCherry+ cells that express TUBB L240F and that of mCherry-cells (uninfected controls) is shown in Fig 3A. The results of this study show that both mCherry+ cells that express TUBBL240F or empty vector and mCherry-cells that represent parental cells are inhibited by rigosertib as well as BI2536 in a concentration-dependent manner with very similar kinetics. Next, we repeated these studies and measured the effect of rigosertib and BI2536 on the viability of K562 cells that express TUBBL240F or empty vector. The results of this study, which are shown in figure 3B, demonstrate that both rigosertib and BI2516 induce cell death in both mCherry+, TUBBL240F-expressing cells and parental (mCherry-) cells with similar kinetics. This effect was also seen with empty vector expressing cells and their parental counterparts. Cells treated with rigosertib at concentrations of 100 or 200nM had less that 10% viable cells on day 6 compared to their untreated controls, suggesting that expression of mutant TUBB had no effect on the growth inhibitory or apoptosis-inducing activities of rigosertib or BI2536.

**Figure 3.**
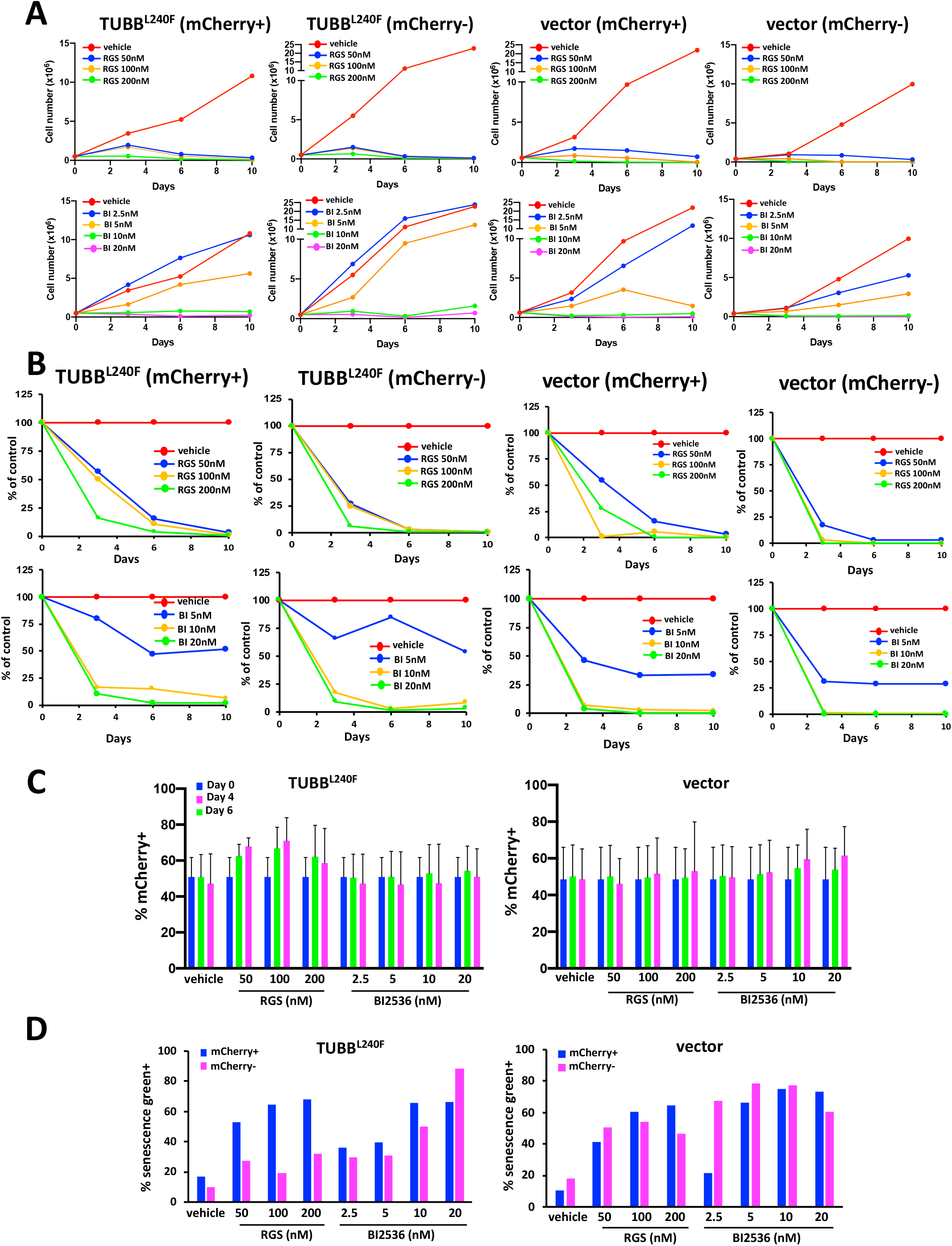
Expression of mutant β-tubulin does not confer a significant survival advantage in K562 cells treated with rigosertib. **(A)** Treatment with rigosertib indices a dose dependent decrease in the proliferation and **(B)** viability of mCherry-positive TUBB^L240F^ and wild-type K562 cells over time. K562 cells were infected with lentiviruses encoding β-tubulin L240F or empty vector and combined with wild-type (mCherry-) cells at a final density of 1.0×10^5^ cells/ml. The cells were then treated with increasing concentrations of rigosertib, BI2536 or vehicle (DMSO) and harvested on days 1, 4 and 6. The percentages of viable (DAPI-), mCherry+ cells were determined using flow cytometric analysis. **(C)** Percent mCherry+ K562 cells expressing TUBB^L240F^ as a function of time. K562 cells were infected with lentiviruses encoding mutant L240F β-tubulin or vector control for 24 hours. The percentage of mCherry+ cells was determined by flow cytometric analysis 48 hrs post-infection and the cells combined in a with wild-type (mCherry-) cells at a final density of 1.0×10^5^ cells/ml. The cells were then treated with increasing concentrations of rigosertib, BI2536 or vehicle (DMSO). Cells were harvested on days 1, 4 and 6 and the percentage of viable (DAPI-), mCherry+ cells determined using flow cytometric analysis. Error bars represent mean+SEM. **(D)** Treatment with rigosertib induces senescence in both wildtype and TUBB^L240F^ K562 cells. K562 cells were infected with lentiviruses as described in A and grown in the presence of increasing concentrations of rigosertib or BI2536 for a 7-day period. mCherry-positive and -negative cells were isolated using fluorescence-activated cell sorting, fixed and the level of β-galactosidase determined by staining with CellEvent senescence green and subsequent flow cytometric analysis.

When we examined the ratio of mCherry+ cells in the small viable fraction that remained at days 4 and 6, we observed a slightly higher fraction of TUBB L240F expressing cells in rigosertib-treated cells (Fig. 3C). Thus, the ratio of mCherry+ cells in vehicle-treated cultures was approximately 50% on days 0,4 and 6 and this ratio of mCherry+ cells increased to 65% to 70% in cells that express L240FTUBB. While this increase was statistically not significant, this trend was repeatedly seen in multiple experiments. This observation is consistent with that made by Jost et al (2017), who interpreted this population to be rigosertib-resistant cells. However, when we examined the surviving cells under the microscope, they were unusually large in size with a cellular morphology that is characteristic of senescent cells. We therefore examined the surviving population for increased levels of β-galactosidase, one of the most reliable markers of senescence. These analyses demonstrated that the majority of the remaining cells in all cultures, including those that express TUBBL250F, exhibited increased β-galactosidase levels as demonstrated by traditional staining (data not shown) as well as flow cytometric analysis using CellEvent senescence green (Fig. 3D), confirming that the remaining cells scored positive for this senescence marker (Fig. 3D). In spite of prolonged incubation in growth media, we were unable to establish cell cultures that can grow in the presence of rigosertib, suggesting that this residual population of cells were truly senescent and do not represent rigosertib-resistant cells.

To further confirm this observation and extend the studies, we developed several stably transfected clonal cell lines that express mutant β-tubulin. We chose to use K562 as well as A549 cells, which are *KRAS* mutant lung carcinoma cells that were shown by us to be sensitive to rigosertib-mediated growth inhibition *in vitro* and *in vivo* (Athuluri-Divakar et al., 2016). For these studies, lentiviral constructs that encoded empty vector or wild-type (wt) *TUBB* or *TUBB* L240F, which were individually transduced into K562 and A549 cells and clones that stably expressed *TUBB* L240 and the mCherry vector selected using limiting dilution. Clones that were >95% mCherry+ by flow cytometric analysis were selected for further study, with a representative K562 and A549 clones shown in figures 4A and 5A (left panels). To determine whether L240F TUBB confers resistance to rigosertib, exponentially growing K562 clones were grown in the presence of increasing concentrations of vehicle or rigosertib for a period of 96 hours. At the end of the study, viability was determined using CellTiter Blue. The results of this study (figure 4A, right panel) show that expression of the L240F beta-tubulin mutant does not confer a significant survival advantage to this cell line.

**Figure 4.**
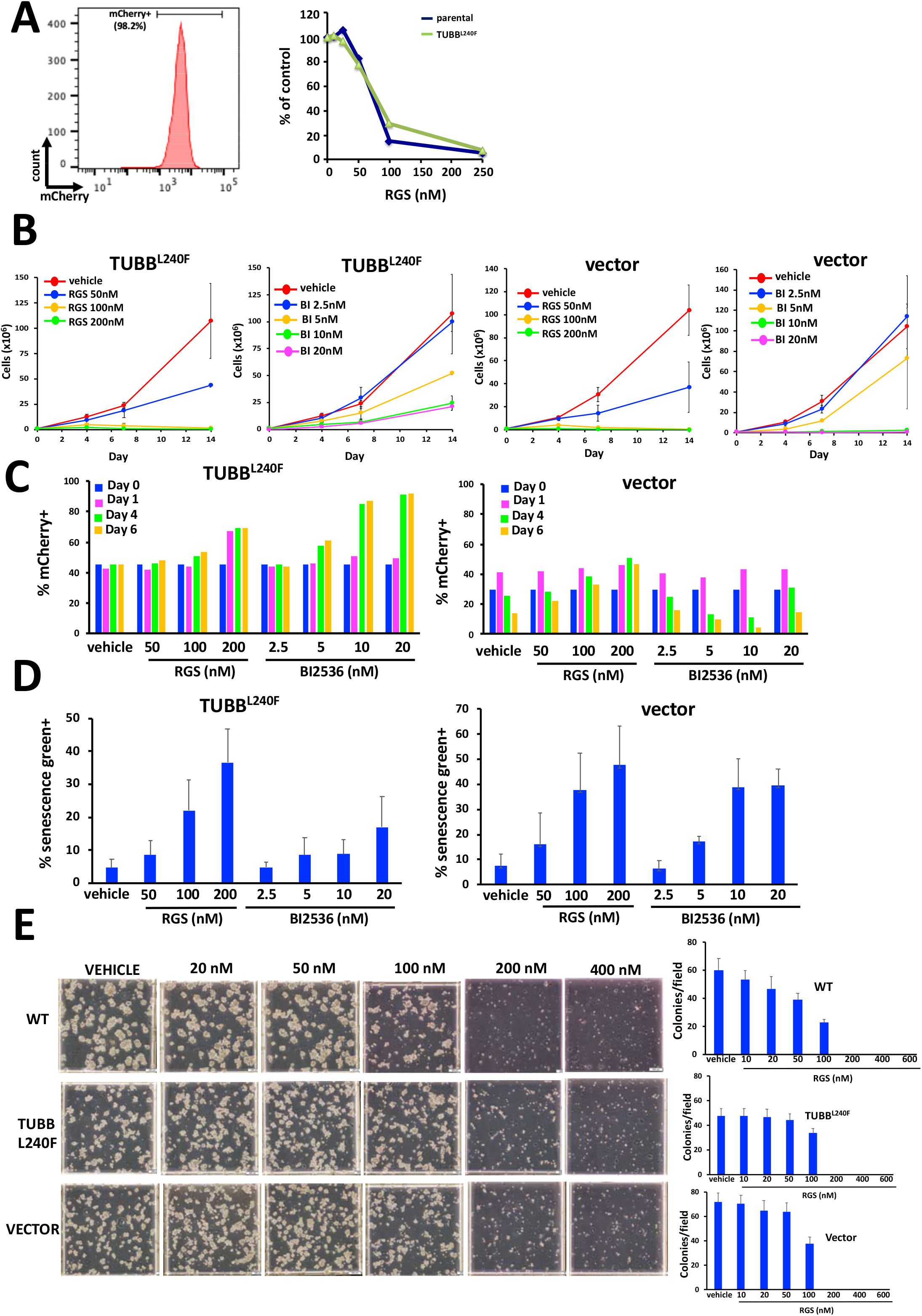
Expression of L240F mutant β-tubulin does not confer resistance to rigosertib. K562 cells were infected with lentivirus that encodes the L240F mutant form of β-tubulin along with the mCherry fluorescence marker and stable cell lines isolated using limiting dilution cloning. **(A)** Flow cytometric analysis of mCherry expression in a representative K562 clone showing that >95% of cells are mCherry+ (left panel). These cells were then treated with the indicated concentrations of rigosertib and the percentage of viable determined at 96 hours post plating. Note that expression of the L240F β-tubulin mutant confers little or no resistance to rigosertib (right panel). **(B)** Growth of L240F β-tubulin expressing K562 cells in the presence of increasing concentrations of rigosertib or BI2536, showing that rigosertib inhibits the proliferation of K562 cells that express TUBB^L240F^. Analysis of one representative clone for each cell line is shown. **(C)** K562 cells expressing TUBB^L240F^ were combined with wt cells and seeded at a density of 1×10^5^ cells/ml. The cells were treated with the indicated concentrations of rigosertib or BI2536 over a 6-day period and the percentage of mCherry+ cells determined by flow cytometric analysis. Note that the percentage of mCherry+ cells also increases in the presence of BI2536, indicating that this is not a rigosertib-specific phenomenon. **(D)** Treatment of K562 cells expressing TUBB^L240F^ or control vector with rigosertib or BI2536 induces senescence. Cells were treated with the indicated concentrations of rigosertib or BI2536 for a 7-day period and the level of β-galactosidase activity in viable cells measured by flow cytometric analysis using CellEvent senescence green. **(E)** 3-dimensional growth of L240F β-tubulin expressing and control K562 cell lines in the presence of increasing concentrations of rigosertib, showing that rigosertib inhibits their proliferation in methylcellulose. Representative images (left panel) and average quantitation of 10 fields per plate in duplicate (right panels) are shown.

To compare the relative sensitivity of TUBBL240F expressing cells with that of parental wt cells, the cells were mixed 1:1 with uninfected cells and treated with DMSO, rigosertib or BI2536. Growth curves of the mCherry+ cells that stably express TUBB L240F and vector control are shown in Fig 4B. Cell cultures expressing L240 mutant tubulin as well as empty vector expressing cells that were treated with rigosertib at concentrations of 100 or 200nM had fewer than 10% viable cells remaining at day 6. When we examined the ratio of mCherry+ cells in this viable fraction, we observed a slightly higher fraction of TUBB L240F expressing cells in rigosertib-treated cultures (Fig. 4C). However, we also observed that this fraction of L240F TUBB cells were increased to a greater degree in BI2536-treated cultures, suggesting that this phenomenon is not unique to rigosertib. When we examined the surviving cells under the microscope, they displayed characteristics of senescent cells such as abnormally large size, with the majority of these surviving cells exhibiting increased β-galactosidase levels as demonstrated by traditional staining (data not shown) as well as flow cytometric analysis using CellEvent senescence green (Fig. 4D).

Next, we examined the growth of wt, TUBBL240F- and empty vector-expressing cells, in the presence of vehicle or rigosertib in methocult, which allows the growth of cells in 3D. The results of this study with parental K562 (WT) and K562 cells expressing TUBBL240F or empty vector is shown in figure 4E. These results show that the growth of all three clonal cell lines is completely inhibited by 200nM rigosertib, further confirming that expression of TUBBL240F does not confer survival advantage to these cells.

Next, we examined the effects of rigosertib and BI2536 on the growth and survival of clonal A549 cell lines that stably expressed L240FTUBB or empty vector. A representative clone of A549 cells that is >95% mCherry-positive is shown in figure 5A (left panel). The results shown in figure 5A (right panel) demonstrate that rigosertib as well as BI2536 inhibit the growth both vector- and L240F mutant tubulin-expressing cells with similar kinetics and in a dose and time dependent manner. Approximately 7-10% of the empty-vector expressing and L240F mutant tubulin expressing cells were viable at the end of 6 days and this viability further decreased to 1-2% at the end of 14 days.

**Figure 5.**
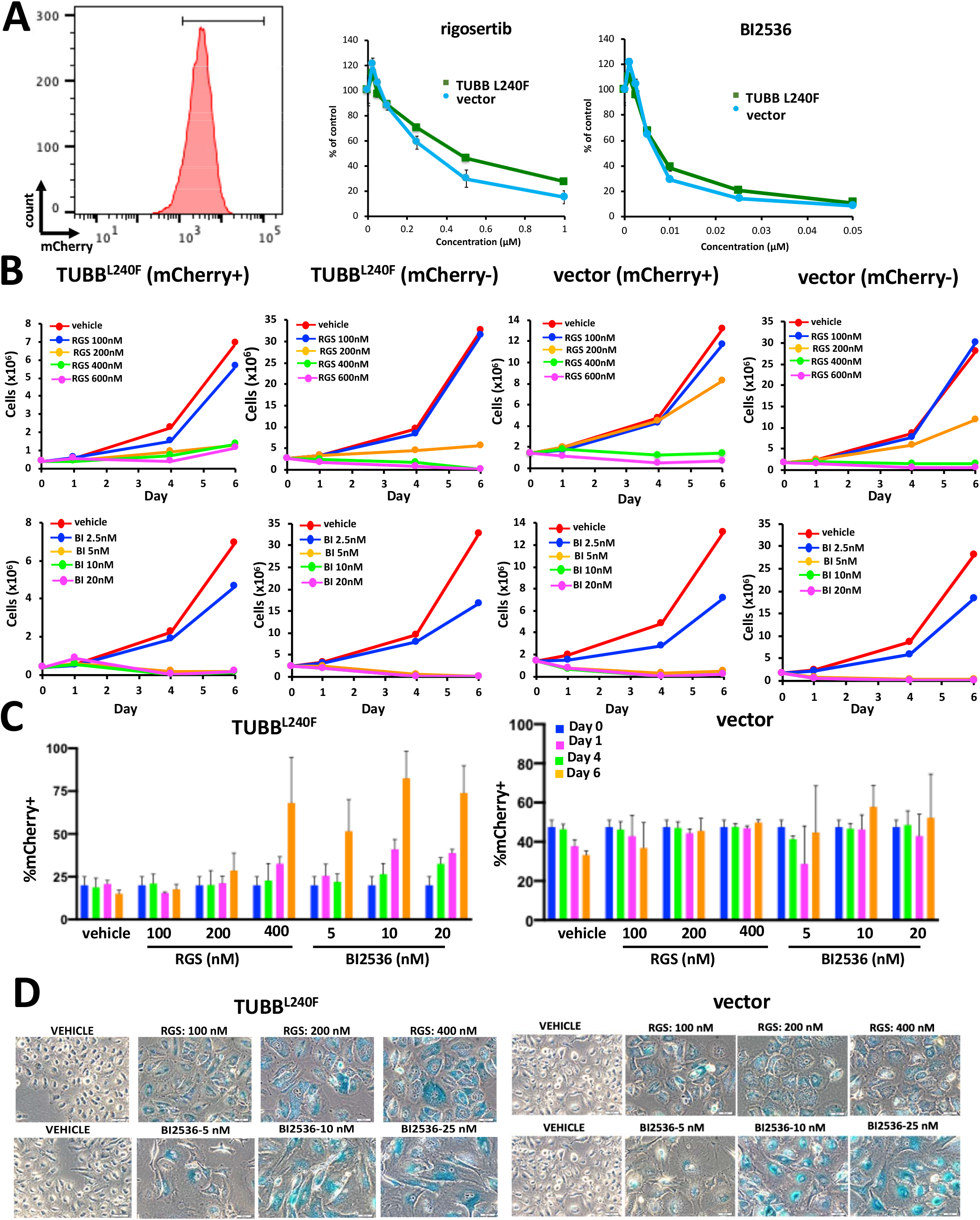
Expression of mutant β-tubulin does not induce resistance to rigosertib in A549 cells. A549 cells were infected with lentivirus that encodes the L240F mutant form of β-tubulin along with the mCherry fluorescence marker. **(A)** Flow cytometric analysis of mCherry expression in representative A549 clones that are >95% of cells are mCherry+ (left panel). The cells were then treated with the indicated concentrations of rigosertib or BI2536 and their viability determined 96 hours post-plating (right panel). **(B)** A549 cells expressing TUBBL240F or vector control were combined with wild-type A549 cells and grown in the presence of increasing concentrations of rigosertib or BI2536 and the growth of mCherrry+ and – cells were measured over a 6-day period. As observed with K562 cells, rigosertib inhibits both the proliferation and survival of A549 cells that express TUBB^L240F^. **(D)** Prolonged treatment with rigosertib and BI2536 induces senescence in A549 cells. A549 cells expressing TUBB L240F or control cells were seeded at a density of 2×10^5^ cells per well in a 6 well dish. The cells were then treated with the indicated concentrations of rigosertib or BI2536 24 hrs post-plating. Media was changed weekly. After 16 days in culture, the cells were washed with PBS, fixed and stained overnight with X-gal (0.1 mg/ml) at 37°C.

To compare the relative sensitivity of TUBBL240F expressing cells with those of wild-type, parental cells, clonal cell lines were mixed with uninfected cells and treated with DMSO, rigosertib or BI2536. Growth curves of the mCherry+ cells that stably express TUBB L240F and that of vector expressing cells are shown in Fig 5B. Cell expressing L240F mutant tubulin as well as empty vector expressing cells that were treated with 400nM rigosertib had fewer than 10% viable cells remaining at day 6. When we examined the ratio of mCherry+ cells in this viable fraction, we observed a slightly higher fraction of TUBB L240F expressing cells in rigosertib-treated cells (Fig. 5C). However, we also observed that the fraction of L240F TUBB cells was also higher in BI2536-treated cultures, confirming that this phenomenon is not unique to rigosertib. Next, we examined the levels of β-galactosidase in the remaining cells to quantitate the percentage of senescent cells in the surviving population. These results, shown in figure 5D, show that 100% of the viable cells treated with 200-400nM rigosertib or 10-20nM BI2536 were senescent (Fig. 5D). This is consistent with published observations that BI2536 is a potent inducer of senescence in certain cell types (Driscoll et al, 2014). As seen with K562 cells, in spite of prolonged incubation in growth media in the presence of rigosertib, we were unable to establish cell cultures that can grow in the presence of this compound, suggesting that this residual population of cells were truly senescent and do not represent rigosertib-resistant cells. These results strongly suggest that the small fraction of cells that survive rigosertib treatment score positive in viability assays that do not distinguish between proliferating and senescent cells and might lead to misinterpretation of drug response.

## DISCUSSION

In an earlier study (Athuluri-Divakar et al, 2016), we provided structural and biochemical evidence to show that rigosertib acts as a RAS-mimetic and binds to the RBDs of the RAF and PI3K family proteins and disrupts their ability to bind to RAS. This conclusion was based on biochemical assays, which included chemical pulldown of target proteins by rigosertib linked to agarose beads, thermal shift assays and NMR spectroscopy. MST showed that rigosertib binds to RAF-RBDs with high affinity, and we were able to solve the structure of the rigosertib-RAF-RBD complex using high resolution NMR spectroscopy. We also demonstrated that this compound inhibits RAS-mediated activation of MAPK and AKT pathways in *in vitro* as well as animal models of RAS-induced tumors (Athuluri-Divakar et al, 2016).

Following the publication of our study, Jost et al (2017) reported that high concentrations of rigosertib (>20μM) exhibited microtubule depolymerizing activity and that exogenous expression of a mutant form of tubulin causes tumors cells to become resistant to the effects of rigosertib. While they did not examine the physical binding of rigosertib to tubulins or RBDs of RAS effectors, they have provided a crystal structure of a molecule in complex with β-tubulin. Rigosertib preparations purchased from a commercial vendor, Selleckchem, were used for these studies. Because commercial preparations of drug candidates are non-GMP quality and are often contaminated with intermediates that are used for the synthesis of the drug and/or degradation products of the final drug candidate, we examined the purity of rigosertib purchased from Selleckchem. Our studies reveal that the preparation examined by us contained approximately 5% impurities and the major impurity seen is ON01500, which is an intermediate in the synthesis of rigosertib. ON01500 is known to be a potent tubulin depolymerizing agent and hence could be the agent that is responsible for the tubulin depolymerization seen by Jost et al (2017). We compared the tubulin depolymerization activities of pharmaceutical grade rigosertib (O-RGS) obtained from Onconova Therapeutics Inc and commercial grade rigosertib purchased from Selleckchem (S-RGS) and ON01500, the intermediate found in S-RGS preparations. Our results show that ON01500 induces complete depolymerization of tubulins at a concentration of 1 μM and that S-RGS induces depolymerization of tubulins at concentrations of 25 μM and above. Since our previous analysis showed that approximately 5% of S-RGS is contaminated with ON01500, the depolymerizing activity seen above 25 μM S-RGS could be entirely attributed to contaminating ON01500. In the case of pharmaceutical grade rigosertib (O-RGS), we did observe a small delay in tubulin depolymerization at concentrations of 50-100μM, which could also be attributed to a small (0.1%) contamination of ON01500. This supposition could also explain why Jost and colleagues also observed approximately a 25% reduction in microtubule growth rates in assays performed using O-RGS at higher concentrations (20μM) (personal communication). Our conclusion that rigosertib does not bind to β-tubulins is further supported by our binding studies using MST (Wienken et al., 2010), where we measured the binding of ON01500, rigosertib and vincristine, a known tubulin depolymerizing agent. These studies clearly demonstrate that while ON01500 and vincristine bind to highly purified preparations of β-tubulin, rigosertib fails to do so.

To determine the crystal structure of rigosertib-tubulin complex, Jost et al (2017) used a concentration of 2mM S-RGS to soak the crystals of β-tubulin. Based on our purity analysis of S-RGS, a 2mM concentration of S-RGS is expected to contain 100μM of ON01500 as an impurity, which could have contributed to the structure provided by Jost et al (2017). ON01500 and rigosertib have minor structural differences and rigosertib is synthesized from ON01500 by converting the amino group of ON01500 into a glycyl moiety. When we examined the electron density for the unique carboxy group of rigosertib in the structure described by Jost et al (2017), we find that the density is very weak and some of the proposed hydrogen bonding interactions are less than optimal. In our opinion, the weak electron density for the carboxy group in the reported structure can be interpreted as ON01500 and a water molecule, particularly in light of our data showing the presence of ON1500 in commercially sourced rigosertib, and the capacity of ON01500 to bind and depolymerize tubulins.

Jost et al (2017) also reported that expression of the L240F β-tubulin mutant provides resistance to rigosertib, indicating that tubulin binding is critical to its cytotoxic activity. For these experiments, Jost et al. prepared lentiviral constructs that encoded empty vector, wt *TUBB* or *TUBB* L240F which were individually transduced into tumor cells. These cell lines, which express the mCherry marker, were combined in a 1:1 ratio with their respective parental lines, treated with rigosertib or DMSO and the fraction of *TUBB*-expressing cells measured up to 7 days after treatment as mCherry-positive cells by flow cytometry.

To assess whether expression of L240F *TUBB* confers resistance to rigosertib, we obtained lentiviral vectors that encode wt tubulin or the L240F beta-tubulin mutant from Dr. Weissman’s laboratory and developed several stably transfected clonal K562 and A549 cell lines that express these tubulin proteins. When exponentially growing A549 and K562 clones were treated with increasing concentrations of vehicle or rigosertib, both cell lines that express wt or mutant L240F β-tubulin underwent apoptosis with similar kinetics, suggesting that stable expression of L240F β-tubulin does not confer resistance to rigosertib.

In their study, Jost et al. seeded mutant tubulin-expressing and wt cells in a 1:1 ratio in the same well and measured the ratio of the two populations after treatment with rigosertib or DMSO by flow cytometry, making use of the fact that the tubulin construct is marked with mCherry. An elevated ratio of mCherry-expressing cells after rigosertib treatment compared to that of DMSO was interpreted to indicate that expression of L240F mutant tubulin confers resistance. To determine the reason for the discrepancy between our studies, we repeated the studies as described by Jost et al using K562 cell line which was infected with empty vector or the TUBB L240F expression vector. Seventy-two hours post-infection, the cells were mixed with uninfected cells and treated with DMSO, rigosertib or BI2536, a pan-PLK inhibitor as a control. The results of this study showed that both wild-type cells and those expressing TUBB L240F underwent growth inhibition and apoptosis in the presence of rigosertib and BI2536 with similar kinetics, suggesting that expression of mutant TUBB had no effect on the growth inhibitory activities of rigosertib or BI2536. Cell cultures treated with rigosertib at concentrations of 150 or 200nM had fewer than 10% viable cells remaining at day 4, and less than 1% at day 14. When we examined the ratio of mCherry+ cells in the residual cell fraction at day 4, we observed a slightly higher ratio of mCherry+ cells in TUBBL240F-expressing cell population. Further analysis demonstrated that the majority of the surviving cells exhibited increased β-galactosidase levels as demonstrated by traditional staining as well as flow cytometric analysis using CellEvent senescence green, confirming that the remaining cells underwent senescence. In spite of prolonged incubation in growth media, we were unable to establish cultures that could grow in the presence of rigosertib at concentrations of 200-300nM, suggesting that this residual population of cells were truly senescent and do not represent rigosertib-resistant cells. Next, we repeated these studies with K562 and A549 cell lines that were stably infected with TUBBL240F expression vectors and selected for mCherry expression. When these cells were mixed with their parental cell lines and examined for the effects of rigosertib or BI2436, we again observed that both TUBBL240F expression did not confer any growth advantage. At the end of 4 days of incubation with rigosertib, both parental cells and their TUBB240F expressing counterparts underwent apoptosis with similar kinetics and by this time, less than 10% of cells in each population were viable. When we examined the ratio of mCherry+ cells in this viable fraction, we observed a slightly higher percentage of TUBB

L240F-expressing cells in rigosertib-treated cells. However, we also observed that this population of L240F TUBB cells was also higher in BI2536-treated cells, suggesting that this phenomenon is not unique to rigosertib. More importantly, when we examined β-galactosidase levels in the surviving cells to quantify the percentage of senescent cells, we found that nearly 100% of the remaining viable cells treated with 200-400nM rigosertib or 10-20nM BI2536 were senescent (figures 4 and 5).

Since a definitive proof of drug resistance is to establish permanent cell lines that grow in 4-5-fold higher concentrations of the drug, we made repeated attempts to establish permanent cell lines that express L240F-TUBB and proliferate in the presence of 3-4-fold higher concentrations of rigosertib which are otherwise lethal to wt cells. Despite several attempts, we have not been able to establish any cell lines that grow at a concentration that is otherwise lethal to wild-type cells. Based on these set of studies, we conclude that L240F mutant form of β-tubulin does not confer resistance to rigosertib.

Rigosertib is a small molecule therapeutic agent currently in Phase III clinical trials for the treatment of myelodysplastic syndrome and cancers that are commonly RAS-mutated. This drug has been administered to more than 1,000 cancer patients and none of these patients experienced the typical side effects that are induced by tubulin depolymerizing agents such as peripheral neuropathy, bone marrow suppression, hair loss and gastrointestinal problems (Lu et al, 2012), further suggesting that rigosertib has a different mechanism of action compared to any of the known tubulin-binding agents.

Based on this analysis, we conclude that some of the results obtained by Jost et al (2017) are due to contamination of commercial grade rigosertib with ON01500, which is an intermediate in the synthesis of rigosertib and is a potent tubulin polymerizing agent. In addition, the assays used to determine resistance relied on short term culture assays which did not take into account that many targeted therapies induce senescence of tumor cells which were scored as proliferating cell population in flow cytometric analyses.

## METHODS

### Chemicals

GMP-grade rigosertib (RGS) and ON01500 were obtained from Onconova Therapeutics, Inc. Chemical grade RGS and BI2536 were purchased from Selleck Chemicals. Vincristine was purchased from Sigma Aldrich.

### Plasmids and Generation of Lentiviruses

Lentiviral supernatant and plasmids encoding wildtype and L240F-mutant *TUBB* were obtained from Jonathan Weissman (ref) and co-transfected with packaging plasmids (Singleton & Wood, 2016) using Lipofectamine 2000 (Thermo Fisher Scientific) into HEK-293T cells to generate recombinant lentiviruses. Viral supernatant was harvested 48 and 72 hrs post-transfection and filtered prior to further use.

### Cell Lines

HEK-293 and A549 cells were cultured in Dulbecco’s Modified Eagle Medium (DMEM) (Life Technologies) supplemented with 10% FBS and penicillin-streptomycin. K562 cells were obtained from Jonathan Weissman and grown in RPMI (Life Technologies) supplemented with 10% FBS and penicillin-streptomycin. All cells were at 37°C grown under humidified conditions and 5% CO_2_.

### Generation of Tubulin-expressing cells

Tubulin-expressing and vector control cell lines were generated by spinoculation. Briefly, 1×10^6^ cells were infected with 100-150μl of virus in the presence of 7.5mg/ml polybrene and centrifuged at 2,200 rpm at 37°C for 1 hr. Media was changed the following day and the cells grown for an additional 48 hrs prior to use. Stable transfectants were infected as described above, grown in the presence of 2μg/ml puromycin and clonal cell lines further selected using limiting dilution. The percentage of mCherry+ cells was determined by flow cytometric analysis as described below. Cell lines with >95% mCherry+ cells were chosen for further analysis.

### Growth and Viability Assays

To determine the growth kinetics of tubulin-expressing K562 and A549 cells, clonal cell lines were seeded at a density of 1×10^5^/ml (K562) or 1×10^6^ cells in a 100mm dish (A549). Cell number and viability determined on the indicated days by trypan blue exclusion. For 96 hr dose response assays, exponentially growing cells were seeded at a density of 2.5×10^3^/well of a 96-well plate and treated the same day (K562) or the following day (A549) with the indicated concentrations of GMP-grade RGS. Viability was determined 96 hours posttreatment using CellTiter Blue (Promega) according to the instructions of the manufacturer.

To assess proliferation in 3-demensional assays, 1×10^5^ cells were seeded in 35mm dishes in duplicate in H4100 base methocult (StemCell Technologies, Inc.), supplemented with 10% FBS and the recommended volume of IMDM in the presence increasing concentrations of rigosertib. Plates were incubated for 6 days at 37°C under humidified conditions and 5% CO_2_. Images were acquired using an Olympus microscope using a 20X objective and manufacturer supplied software.

### Tubulin Polymerization Assays

25μg of MAP-rich tubulin (Cytoskeleton, Inc.; catalog # ML116) in general tubulin buffer (80mM PIPES, pH 6.9/ 2mM MgCl_2_/ 0.5mM EGTA/10μM fluorescence reporter) (catalog # BP01; Cytoskeleton, Inc.) was combined with 1mM GTP, 15% tubulin glycerol cushion buffer (Cytoskeleton, Inc.; catalog # BPST05-001) and the indicated concentrations of rigosertib (obtained from either Selleckchem [chemical grade] or Onconova Therapeutics, Inc. [GMP/pharmaceutical grade]), ON01500 (Reddy et al., 2011), nocodazole (Sigma-Aldrich; catalog # M1404) and vincristine (Sigma-Aldrich; catalog # V8388). All compounds were dissolved in DMSO and used as 25X stock solutions. The reactions were incubated at 37°C in a Synergy H1 Multi-mode Hybrid Reader (BioTek) and fluorescence measured every 30 seconds over the indicated period of time. Fluorescence values were normalized by subtracting the points obtained at the start of polymerization for each reaction from all subsequent data points.

### Microscale Thermophoresis

Tubulin (Cytoskeleton, Inc.; catalog # T240) was labeled for microscale thermophoresis (MST) using the Monolith NT Protein RED-NHS Labeling Kit (NanoTemper Technologies, München, Germany) according to the instructions of the manufacturer. Briefly, tubulin at a concentration of 20μM was incubated with 2X dye at a ratio of 1:1 in labeling buffer (manufacturer supplied) in the dark at room temperature for 30 minutes. Free dye was removed using a manufacturer supplied gel filtration column and the protein eluted in 0.5ml of general tubulin buffer (80mM PIPES, pH 6.9/ 2mM MgCl2/ 0.5mM EGTA). To determine the Kd values of rigosertib, ON01500 and vincristine to tubulin, labeled tubulin was incubated with increasing concentrations of compound in reaction buffer (20mM Tris-HCl pH 7.5/ 150mM NaCl/ 1mM MgCl_2_/ 2% DMSO/ 0.5% prionex) for 30 minutes in the dark at room temperature. All compounds were used as 100X stock solutions. The samples were then centrifuged at 13,000 rpm for 2 minutes before being loaded into standard capillaries provided by the manufacturer. Fluorescence values from the binding reactions were determined using the Monolith NT.115 (NanoTemper Technologies, München, Germany). Binding data was analyzed using GraphPad Prism (GraphPad Software, San Diego, CA) to determine K_d_ values. For these analyses, the fluorescence value from the thermophoresis plots corresponding to the lowest concentration of compound used in the titration was subtracted from every data point prior to normalization. In the case of rigosertib (for which a curve could not be fit), the highest value was set to 100 and the data normalized accordingly.

### Flow Cytometry

For analysis of mCherry expression and sorting of viable mCherry+/− cells, single cell suspensions were prepared in PBS supplemented with 2% FBS and 1μg/ml DAPI. Levels of senescence in K562 clonal cell lines were measured using the CellEvent Senescence Green flow cytometry assay kit according to the instructions of the manufacturer (Thermo Fisher Scientific) except that cells were stained with fixable viability dye eFluor 450 (Thermo Fisher Scientific) for 30 min at 4°C prior to fixation according to the manufacturer’s instructions. To analyze β-galactosidase levels in cultures that were a mixture of mCherry+ and wild-type (mCherry-) cells, viable (DAPI-) mCherry+ and mCherry-cells were sort purified prior to fixation in 2% paraformaldehyde, washed extensively with phenol red-free RPMI containing 2% FBS and stored overnight at 4°C prior to staining with CellEvent Senescence Green (Thermo Fisher Scientific). Flow cytometric analysis and sorting was performed at the Flow Cytometry shared Resource Facility at the Icahn School of Medicine at Mount Sinai. Data for analytical purposes were acquired using an LSRFortessa X-20 or FACSCantoII (BD Biosciences). Sorted cell populations were obtained using an Arial II (BD Biosciences) high-speed sorter. All data were analyzed using FlowJo v10 (Treestar) software.

### Senescence Studies

β-galactosidase levels in K562 cell lines were determined as described in the flow cytometry section. To analyze levels of β-galactosidase in A549 cells, cells were washed with PBS and fixed in PBS containing 2% formaldehyde/0.2%gluteraldehyde for 5 minutes at room temperature. The cells were then washed twice with PBS and stained with 0.1mg/ml X-gal/40mM citric acid sodium phosphate buffer (0.1M citric acid monohydrate/0.2M Na_2_HPO_4_), pH 6.0/ 5mM potassium ferrocyanide/ 5mM ferricyanide/ 150mM NaCl/ 2mM MgCl_2_ overnight at 37°C. Images were acquired using an Olympus microscope using a 20X objective and manufacturer supplied software.

### Statistical Analysis

Statistical analysis was performed using a standard, unpaired, two-tailed Student t test. Data are graphed as mean ± SEM.

## AUTHOR CONTRIBUTIONS

SJB performed the tubulin polymerization assays, MST, cell-based assays and flow cytometric studies. SCC generated cell lines and performed cell-based assays. SKAD and MVRR performed the tubulin polymerization assays and MST. RV and RJ performed the structural studies. AA and EPR designed studies, analyzed data and wrote the paper.

## ACKNOWLEDGEMENTS

We thank Dr. Jonathan Weissman for providing us with lentiviral constructs encoding wild-type and L240F tubulin as well as lentiviruses encoding the same. This work was supported by grants from Onconova Therapeutics Inc. (Newtown, PA), the U.S. Army Medical Research and Materiel Command (LC160287) and the National Institutes of Health (NIH) (5R21CA227963-02) to EPR. Use of the flow cytometry shared resource facility was supported by a NIH Cancer Center Support Grant (P30CA196521) to the Tisch Cancer Institute. EPR is an equity holder, board member and a paid consultant of Onconova Therapeutics, Inc. SJB is a paid consultant of Onconova Therapeutics Inc. MVRR and SCC are stockholders and paid consultant of Onconova Therapeutics Inc. SJB, SCC, MVRR and EPR are named inventors on pending and/or issued patents filed by Temple University.

